# Inter-group Heterogeneity of Regional Homogeneity (REHO)

**DOI:** 10.1101/2022.08.29.505722

**Authors:** Yan Jiang, Mohammed Ayoub Alaoui Mhamdi, Russell Butler

## Abstract

Regional Homogeneity (REHO) measures the similarity between the time series of a given voxel and those of its neighbors. First discovered in a task-activation paradigm, REHO was considered as a complementary method to model-driven analysis of fMRI time series. With the increased popularity of resting-state paradigms, REHO has become a widely used method for inferring neural activity in the resting state. However, the neural/physiological processes that give rise to REHO are poorly understood. Differences in REHO across groups may not be indicative of differences in neuronal activity. Here, we investigate physiological contributions to REHO across 412 subjects in 9 separate datasets downloaded from OpenNeuro where both physiological signals (respiratory rate, heart rate, and motion) and resting state data are available. Overall, we find an inverse correlation between heart rate and REHO across subjects, an inverse correlation between respiratory rate and REHO across time, and differences in REHO across groups is driven primarily by FWHM of data and motion. We conclude that, due to REHO’s highly significant correlation with motion, heart rate, and respiratory rate, REHO should be used with caution to infer differences in neuronal activity across groups.

## 1 Introduction

Regional homogeneity (REHO) is a data-driven method for analyzing functional magnetic resonance imaging(fMRI) data. REHO originates from Kendall’s coefficient of concordance (KCC) (Kendall and Gibbons, 1990), which was used to measure similarity between the time series of a given voxel to those of its nearest neighbors. Due to the fact that fMRI signal is a relative hemodynamic change linked to neuronal activity (Butler et al., 2017, 2019, 2020; Cote et al., 2021; Forouhandehpour et al., 2021), REHO is able to detect unpredicted hemodynamic responses that model-driven method failed to find and has been used to understand the high complexity of the human brain function (Zang et al., 2004). Unlike model-driven methods REHO is robust to variability across trials (Zang et al., 2004), which makes REHO a promising method for within group and between groups analyses. Resting state fMRI (rs-fMRI) is easy to implement because it does not require the subjects to perform a task. Rs-fMRI can provide more functional information, helping to better understand the mechanisms underlying specific diseases (Shao et al., 2019). Analysis of rs-fMRI, however, is susceptible to confounding factors, due to the lack of strong stimulus-driven response. REHO in the Default Mode Network (DMN) has been observed to increase during rest (Long et al., 2008) and decrease during task engagement Zang et al. (2004). As a result, REHO is a promising method of the study of resting-state fMRI and has been used for a large number of studies on the diseased brain. Several studies have shown significant REHO differences observed in patients compared to healthy controls. Below we list the disorders followed by the brain area in which a difference was found. In recent researches, increased or decreased REHO in certain brain regions has been associated with Methamphetamine Dependence (Yu et al., 2020), Parkinson’s Disease (Liu et al., 2019b; Li et al., 2020), Drug-Naive Bipolar Disorder (Shan et al., 2020), Schizophrenia (Ji et al., 2020), psychiatric disorders (Wei et al., 2020), seizures (Liu et al., 2019a), obsessive-compulsive disorder (Yang et al., 2019), manic pediatric bipolar disorder (Xiao et al., 2019), white matter hyperintensities and cognitive impairment (Ye et al., 2019), Strabismus and Amblyopia (Shao et al., 2019) and corneal ulcer (Xu et al., 2019), psychiatric disorders (Wei et al., 2020), Alzheimer (Cheng et al., 2019), Vascular mild cognitive impairment (Zuo et al., 2019). Figure 1 demonstrate brain regions where REHO is most associated to mental disorders or other diseases mentioned in recent REHO researches. We counted how often each region appeared in those studies shown in figure 1. Frontal gyrus, angular gyrus, occipital gyrus, cingulate gyrus and cerebellum cortex had been mentioned the most over 4 times. All of these researches performed motion corrections and some of them had no band pass preprocess. Only half of them performed nuisance regression and none of them performed smoothing to certain FWHM on fMRI data.

**Fig. 1.**
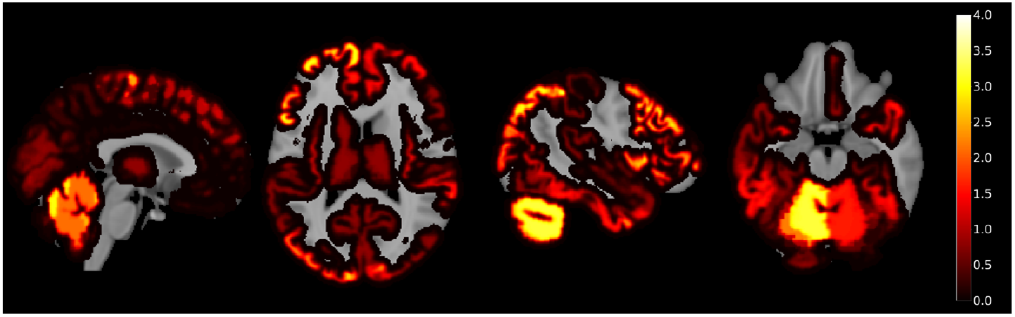
Maps of regions associated with diseases and number of times appeared in recent researches.

While these studies use REHO as a proxy for increases/decreases in neuronal activity, the blood oxygenlevel-dependent (BOLD) signals, which rs-fMRI measures, are observed as being severely affected by multiple non-neuronal artifacts such as cardiac rate, respiratory rate and head motion. BOLD signal changes are correlated with respiratory rate due to the respiration-induced modulation of blood flow and CO_2_ in the brain. Subtle variations in breathing depth and rate that occur naturally during rest can therefore account for a significant amount of variance in the BOLD signal (Chang et al., 2009). Changes in cardiac rate, which could affect both blood volume and flow to the brain, also show significant correlations with BOLD signals (Shmueli et al., 2007). The significant correspondence between the global average brain signal and both the respiratory rate and cardiac rate indicates that much of the common signal expressed by voxels is due to these physiological processes (Chang et al., 2009). In addition, head motion-induced artifacts contribute substantially to the rs-fMRI signal, and produce systematic but spurious patterns in correlation. The relationship between head motion and changes in the BOLD signal remains even after data realignment and regression of realignment estimates and their derivatives from the data (Power et al., 2011). While removal of cardiac rate, respiratory rate and head motion has become a standard part of fMRI pipelines, residual contamination may still result in systematic effects that can bias analysis results. While the influence of physiology and motion on rs-fMRI is widely acknowledged, the effects of those non-neuronal artifacts on REHO has not been adequately studied. In this study, we focus on identifying the influences of cardiac rate, respiratory rate and head motion on REHO, to demonstrate the necessity of dealing with those artifacts when using REHO to quantify differences in brain activity across individuals or groups.

## 2 Materials and Methods

### 2.1 Datasets

Our study includes 9 separate datasets from Openneuro.org where both physiological signals (respiratory rate, cardiac rate, and motion) and resting state BOLD data are available. Overall, there are resting-state fMRI with physiological data of 412 subjects including 352 healthy subjects and 60 with diseases. They are 167 men and 241 women with average age 25.3. All fMRI data are re-sampled to the same TR=2s and registered to the same voxel size 3mm x 3mm x 4mm as one subject of dataset UCLA_control. Table 1 contains the details of 9 datasets.

**Table 1.**
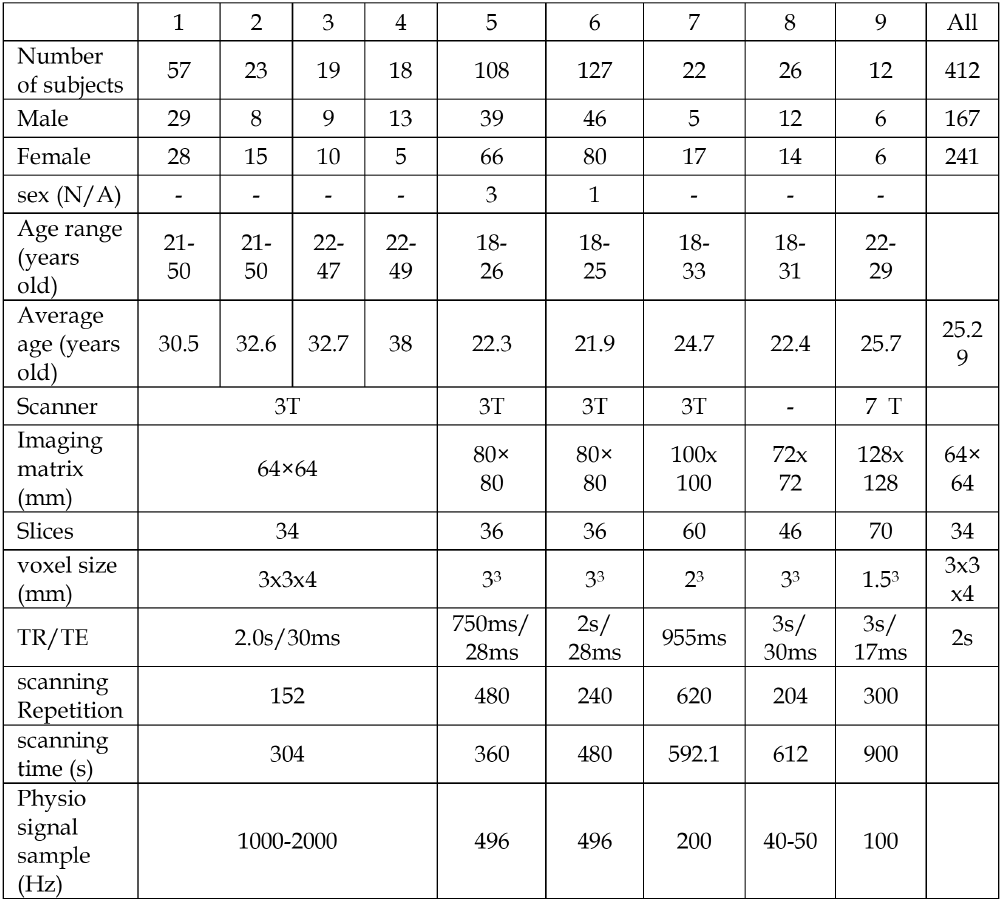
Datasets description. 1: High-resolution-7T; 2: UCLA-control; 3: UCLA-adhd; 4: UCLA-bipolar; 5: UCLA-schz; 6: AOMIC-PIOP1; 7: AOMIC-PIOP2; 8:InterTVA; 9:Multi-echo_fMRI.

#### 2.1.1 UCLA Consortium for Neuropsychiatric Phenomics LA5c Study (UCLA_control, UCLA_adhd, UCLA_bipolar, UCLA_schz)

Four data sets were acquired from Openneuro.org under the accession number ds000030, revision 1.0.0. (Bilder et al., 2020; Poldrack et al., 2016). The subject population consists of healthy controls (130 subjects), as well as participants with diagnoses of adult ADHD (43 subjects), bipolar disorder (49 subjects) and schizophrenia (50 subjects). Due to missing data and bad quality of physiological recording data, only data from 57 subjects of healthy controls, 23 subjects with adult ADHD, 19 subjects with bipolar disorder and 18 subjects with schizophrenia were included in the study. During the resting state scans, Participants were asked to remain relaxed and keep their eyes open; they were not presented any stimuli or asked to respond during the scan. Physiological data were collected using a BIOPAC MP150 with Pulse Oximeter and Respiration modules. Data were sampled at 1000 Hz and recorded using BIOPAC Acknowledge software.physiological recordings were converted from raw data (Acknowledge format, BioPac) using the Bioread python package (https://github.com/njvack/bioread).

#### 2.1.2 Amsterdam Open MRI Collection (AOMIC_PIOP1)

The data set, AOMIC-PIOP1, was acquired from Openneuro.org under the accession number ds002785, revision 2.0.0. (Snoek et al., 2020b,a). The subject population consists of healthy participants (216 subjects). Due to missing data and bad quality of physiological recording data, only data from 108 subjects were included in the study. During the resting state scans, participants were instructed to keep their gaze fixated on a fixation cross in the middle of the screen with a gray background (RGB: [150, 150, 150]) and to let their thoughts run freely. Respiratory traces were recorded using a respiratory belt (air filled cushion) bound on top of the subject’s diaphragm using a velcro band. Cardiac traces were recorded using a plethysmograph attached to the subject’s left ring finger. Data was transferred to the scanner PC as plain-text files (Philips “SCANPHYSLOG” files) using a wireless recorder with a sampling frequency of 496 Hz. Physiology files were converted to BIDS-compatible compressed TSV files using the scanphyslog2bids package.

#### 2.1.3 Amsterdam Open MRI Collection (AOMIC_PIOP2)

The data set, AOMIC-PIOP2, was acquired from Openneuro.org under the accession number ds002790, revision 2.0.0. (Snoek et al., 2020b). The subject population consists of healthy participants (226 subjects). Due to missing data and bad quality of physiological recording data, only data from 127 subjects were included in the study. Other information is the same as the data set AOMIC_PIOP1.

#### 2.1.4 A high resolution 7-Tesla resting-state fMRI test-retest dataset with cognitive and physiological measures (High-resolution-7T)

The data set, High-resolution-7T, was acquired from Openneuro.org under the accession number ds001168, revision 1.0.1. (Gorgolewski et al., 2015). The subject population consists of healthy participants (22 subjects). Due to missing data and bad quality of physiological recording data, only data from 12 subjects of the first run of the second session were included in the study. During the resting state scans, participants were instructed to stay awake, keep their eyes open and focus on a cross. During the scan the participants’ pulse was monitored using a pulse oximeter. Breathing was measured using a pneumatic sensor. Both breathing and pulse signals were recorded using Biopac MP150 system (Biopac Systems Inc., USA) sampled at 5,000 Hz with the Biopac Acqknowledge 4.1 software (Biopac Systems Inc., USA). Physiological data were down-sampled to 100 Hz.

#### 2.1.5 InterTVA. A multimodal MRI dataset for the study of inter-individual differences in voice perception and identification (InterTVA)

The data set, InterTVA, was acquired from Openneuro.org under the accession number ds001771, revision 1.0.2. (Aglieri et al., 2019). The subject population consists of healthy participants (39 subjects). Due to missing data and bad quality of physiological recording data, only data from 22 subjects of the first run of the second session were included in the study.

#### 2.1.6 Multi-echo fMRI replication sample of autobiographical memory, prospection and theory of mind reasoning tasks (Multi-echo_fMRI)

The data set, Multi-echo_fMRI, was acquired from Openneuro.org under the accession number ds000210, revision 00002. (DuPre et al., 2018). The subject population consists of healthy participants (31 subjects). Due to missing data and bad quality of physiological recording data, only data from 26 subjects of the first run of the second session were included in the study.

### 2.2 Data processing

#### 2.2.1 Repetition time (TR) resampling

As we investigate motion effects on group basis, we should keep all dataset in the same TR to reduce the impact of different TR on motion parameter estimation. Datasets UCLA-control, UCLA-adhd, UCLA-bipolar, UCLA-schz and AOMIC-PIOP2 already have the same TR=2s, we applied AFNI command 3dUpsample on dataset the other datasets AOMIC-PIOP1, InterTVA, Multi-echo_fMRI and high-resolution-7T to make sure that all datasets have same repetition time. For the datasets Multi-echo_fMRI and highresolution-7T which had original *TR* = 3*s*, we set the resample command parameter as 6 to resample the fMRI data into *TR* = 0.5*s* and then use AFNI command 3dTcat to extract one time point for every 4 to obtain final *TR* = 2*s*. For the dataset AOMIC-PIOP1 which had original TR=0.75s, we set the resample command parameter as 3 to resample the fMRI data into TR=0.25s and then use AFNI command 3dTcat to extract one time point for every 8 to obtain final *TR* = 2*s*. But because the command only accepts integer parameters, for the dataset InterTVA which had original *TR* = 0.955*s*, we only can set the resample command parameter as 2 to resample the fMRI data into *TR* = 0.477*s* and then use AFNI command 3dTcat to extract one time point for every 4 to obtain final *TR* = 1.908*s*. As a result, all datasets were resampled to exact *TR* = 2*s* except for dataset InterTVA which was re-sampled to *TR* = 1.908*s*.

#### 2.2.2 Motion correction

To align and correct the images for slice timing differences, each 3D volume from each data set was registered to its first volume by AFNI command 3dvolreg. 6 motion parameters including roll rotation, pitch rotation, yaw rotation, displacement in the superior direction, displacement in the left direction and displacement in the posterior direction were acquired at the same time from the output of 3dvolreg.

#### 2.2.3 Despiking

AFNI command 3dDespike was applied. It removes spikes from the 3D+time input dataset and writes a new dataset with the spike values are replaced to fit a smooth curve *f* (*t*)

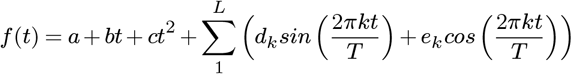

where *T* = duration of time series; The *a, b, c, d, e* parameters are chosen to minimize the sum over *t* of | *v*(*t*) − *f* (*t*) | (L1 regression); *v*(*t*) is voxel time series; The default value of *L* is *NT/*30, where *NT* = number of time points. For each voxel value *v*(*t*), define *s* = (*v*(*t*) − *f* (*t*))*/δ*. Values with *s > c*_1_ are replaced with a value that yields a modified

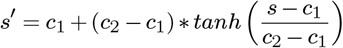

where *c*_1_ = 2.5 (default) is the threshold value of s for a ‘spike’. *c*_2_ = 4 (default) is the upper range of the allowed deviation from the curve: *s* = [*c*_1_.. *∞*) is mapped to *s*^*1*^ = [*c*_1_..*c*_2_).

#### 2.2.4 Registration

Each subject was registered to a template subject (subject 53 of UCLA_control) using FSL’s flirt command. Using the matrix output from this registration, the REHO image of each subject of each dataset was brought into this template subject space. Then, using the matrix output from FSL’s epi_reg (which uses a white matter segmentation boundary to align BOLD images with T1 images), all images were registered to T1-weighted image of subject 53 from UCLA_control.

#### 2.2.5 Bandpass filtering

Data was bandpass filtered from 0.005 Hz to 0.1 Hz using AFNI command ‘3dTproject -passband fbot ftop’. The command removes all frequencies except those in the range between fbot and ftop. We used a conservative high pass value (0.005 Hz) which is slightly lower than conventional the typical value of 0.01 Hz.

#### 2.2.6 Spatial Smoothing

All datasets were smoothed to 8 mm FWHM by using AFNI command ‘3dBlurToFWHM -FWHM 8’. To investigate the effects of smoothing, we performed some analysis pre/post 3dBlurToFWHM (indicated in results).

#### 2.2.7 REHO

AFNI command 3dREHO was applied. Kendall’s *w* (also known as KCC) per voxel was calculated from 3D+time dataset (Zang et al., 2004) described as:

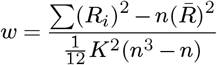

where *w* is the Kendall’s *w* among given voxels; *R*_*i*_ is the sum rank of the ith time point; where 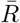 is the mean of the *R*_*i*_’s; *K* is the number of time series within a measured cluster (here, *K* = 27, respectively; one given voxel plus the number of its neighbors); *n* is the number of ranks (here *n* = time points of each 3D+time dataset). Instead of the time series values themselves, Kendall’s *w* can be used to evaluate the similarity of the time series within a cluster of a given voxel and its nearest neighbors. *w* is in range 0-1, with 0 reflecting no trend of agreement between time series and 1 reflecting perfect agreement.

#### 2.2.8 Physiological signals processing

##### Cardiac signal

heart rate was calculated by dividing number of peaks in the cardiac signal by the scanning time. Peaks were determined by python function find_peaks provided by Scipy.org, which uses a wavelet-based method for peak detection.

##### Respiratory signal

respiratory signal of each subject was divided into 10 parts and 10 respiratory rate values (number of peaks divided by scanning time) were defined for each subject. For each dataset, there are (10*numbers of subjects) timepoints in its respiratory series. Figure 2 shows the peaks finding in heart rate raw signals and respiratory raw signals. Figure 3 shows the feature value of respiratory and heart rates of 57 subjects in dataset UCLA-control.

**Fig. 2.**
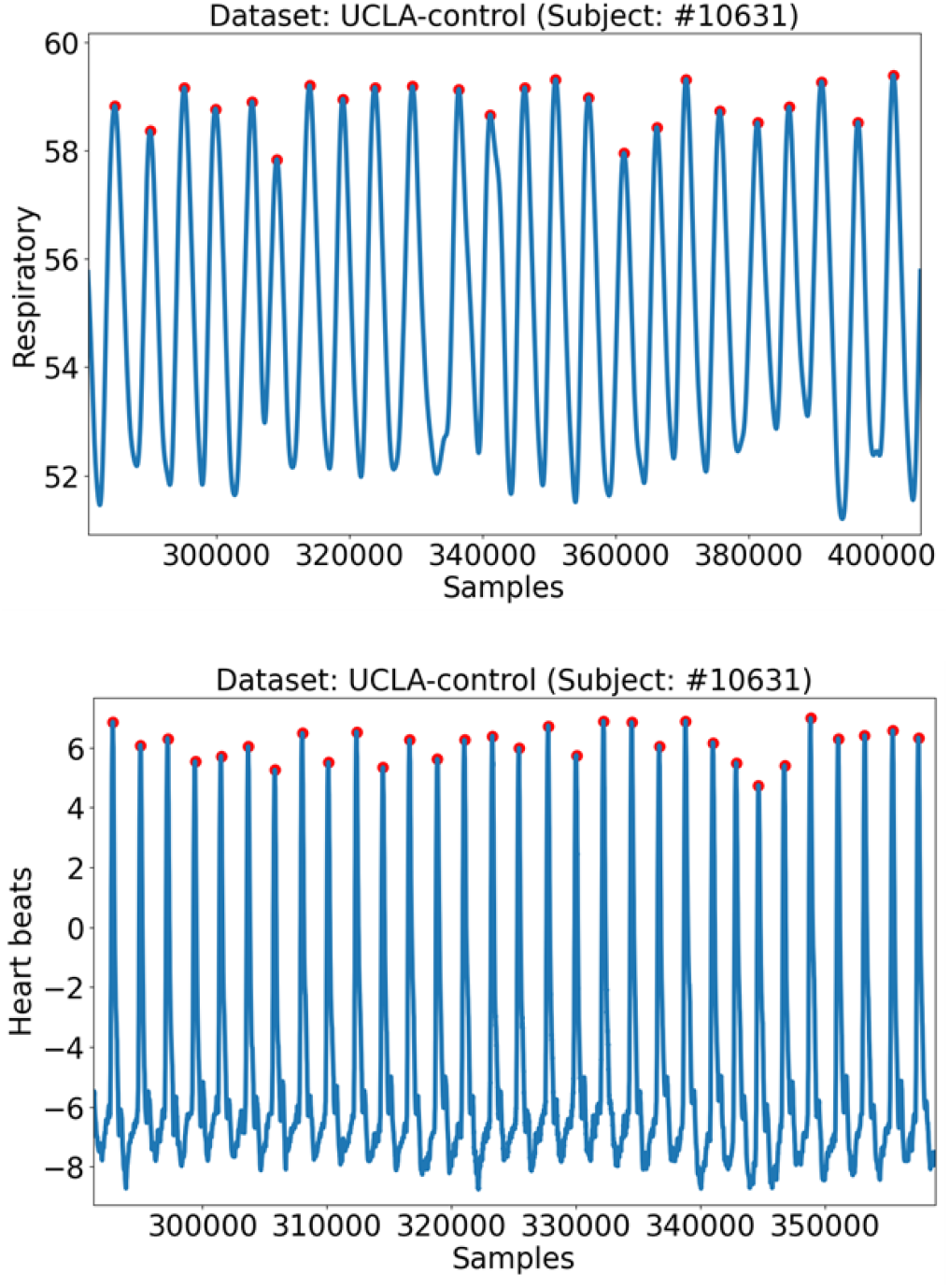
Heart rate, respiratory rate and motion feature values. Respiratory rate signal and Cardiac signal of one subject of dataset UCLA-control. Each red dot presents each peak.

**Fig. 3.**
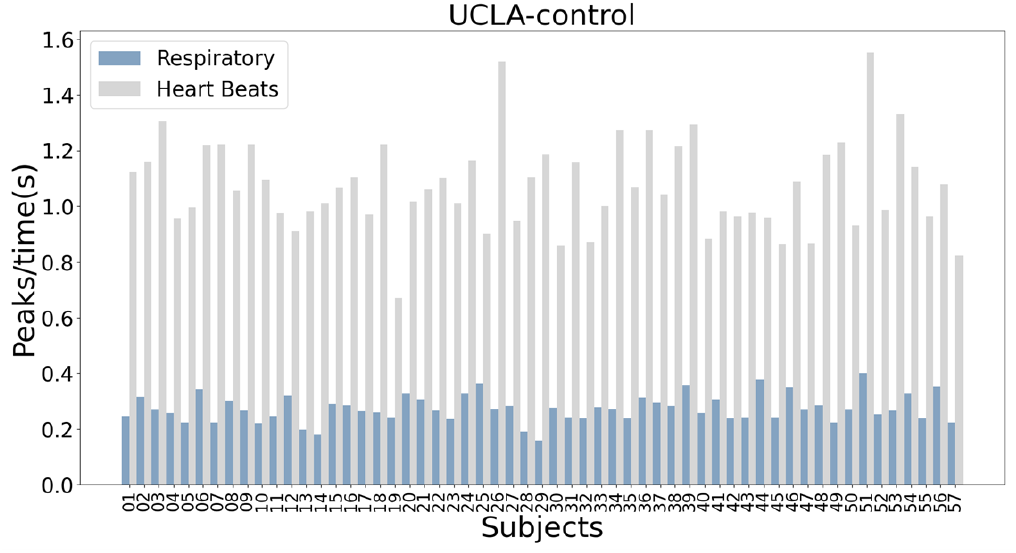
Heart rate, respiratory rate and motion feature values. Feature (peaks/scan time) of respiratory rate and cardiac data of 57 subjects of dataset UCLA-control.)

##### Motion parameters

Motion time series were obtained from fMRI realignment parameters which were acquired by AFNI command 3dvolreg. Realignment parameters represent absolute shifts in position relative to the first volume acquired in a given session and can be used to calculate a relative motion measure between consecutive scans. 6 realignment parameters (roll, pitch, yaw, superior, left and posterior) of all dimensions can be combined to a single normalized motion measure mp. The method described as:

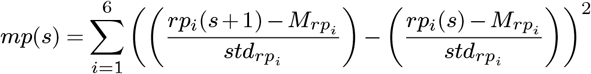

where *rp*_*i*_, *i* denote 6 motion parameters, displacement in the superior direction, displacement in the left direction and displacement in the posterior direction, roll rotation, pitch rotation and yaw rotation, of every *MR* volume relative to the first MR volume is estimated. *M* and *std* denote the respective mean and standard deviations of the realignment parameter (Fellner et al., 2016). Figure 4 shows the average motion parameter of 57 subjects in dataset UCLA-control.

**Fig. 4.**
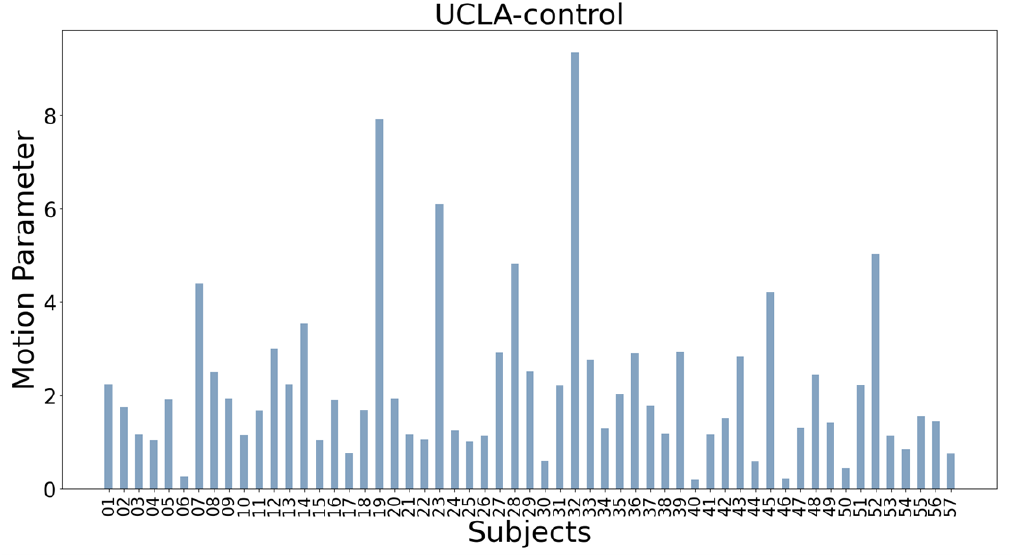
Heart rate, respiratory rate and motion feature values. Single motion parameter combined from 6 realignment parameters of all dimensions (roll, pitch, yaw, superior, left and posterior

#### 2.2.9 Nuisance regression

AFNI command ‘3dTproject -ort f.1D’ was used for nuisance regression. The command runs through all time points and removes the mean of each column in f.1D from data. Average REHO of each REHO image of all 412 subjects are pooled together as the regression target. Nuisances were respiratory rate, cardiac rate and normalized motion measure *mp*. Figure 5 shows the correlation between average REHO and physiological feature time series in different stage of regression (across all subjects). We should note that our nuisance regression was performed at the group level, after calculating REHO (we did not regress directly from the time series).

**Fig. 5.**
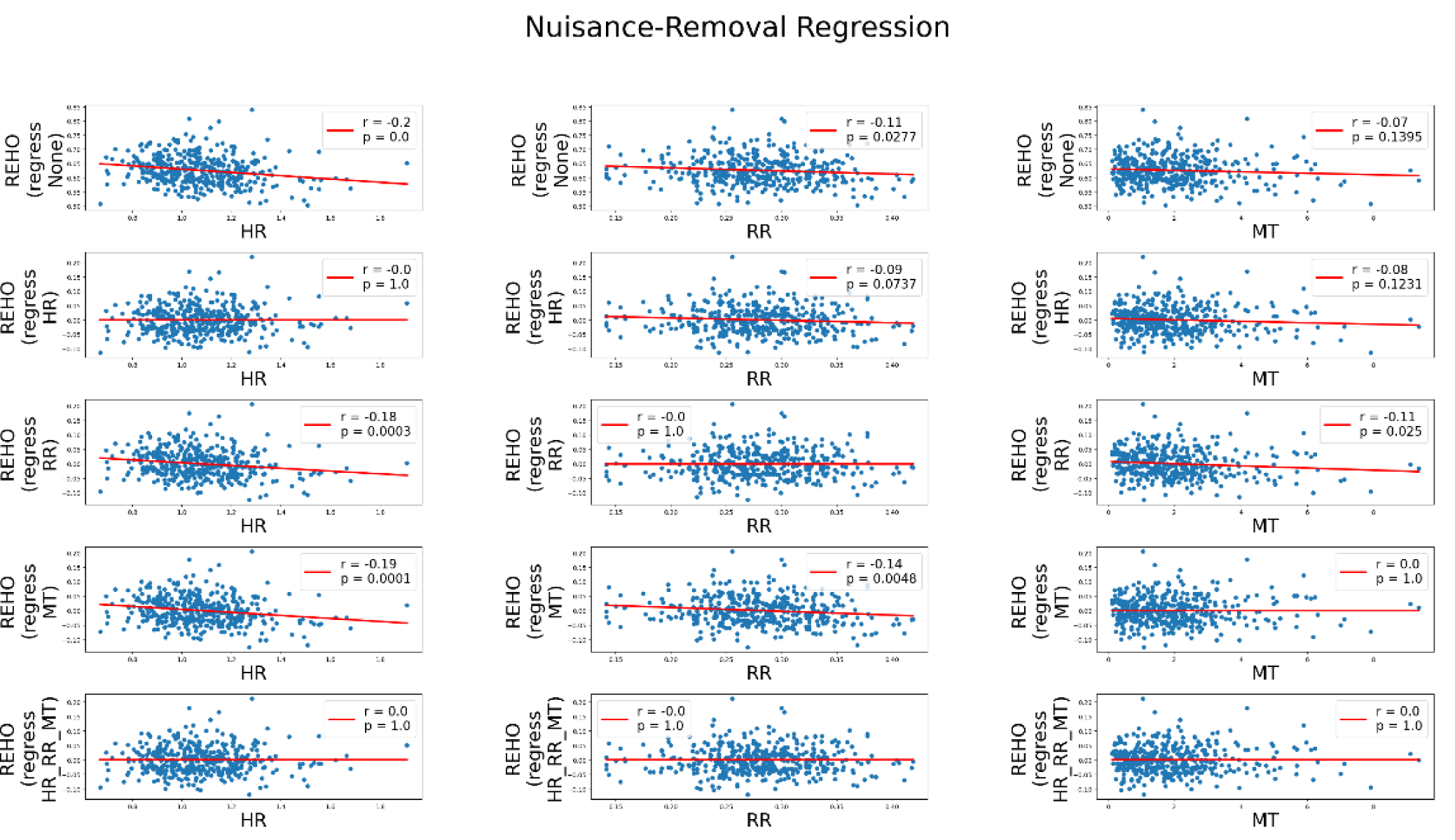
The correlation between average REHO and physiological feature time series in different stage of regression (across all subjects). Without regression (top line), with regression out Heart Rate (HR) only (second line), with regression out Respiratory Rate (RR) only, with regression out Motion only (MT) (third line) and with regression out Heart Rate, Respiratory Rate and Motion.

#### 2.2.10 Statistical analysis

##### Correlation Analysis Heart rate and motion vs REHO

The correlations between REHO data and physiological data were calculated with Pearson’s correlation coefficient (PCC). For each dataset separately, the inter-subject correlation between REHO in each voxel and cardiac rate series was calculated. inter-subject correlation across all 412 subjects (pooling all datasets together) was also calculated. Corresponding p-value was calculated. Correlation with p-value<0.05 is considered as significant (we did not correct for multiple comparisons or family-wise error rate). The calculation of the correlation between REHO and motion was identical to the above (except cardiac rate was replaced with motion parameter mp).

##### Correlation Analysis Respiratory rate vs REHO (dynamic analysis)

For each subject, 3D+time data was divided into 10 segments (across time) and REHO of each individual segment was obtained. Similarly, respiratory signal was divided into 10 segments and respiratory rate was calculated for each segment separately. Each subject then had 10 REHO images and 10 respiratory rate values. Correlations between the 10 REHO images and the respiratory series were calculated in each voxel, for each subject (giving a single correlation map per subject). To calculate pooled dynamic correlations, all 4120 REHO images (total 412 subjects * 10) were combined into a 4D image with 4120 volumes, and respiratory rate vectors were similarly concatenated. The correlation between the 4D REHO and pooled respiratory series was calculated. P-value<0.05 is considered as significant.

##### T-test

Scipy function ttest_ind was applied to investigate significance of differences between average heart rate / respiratory rate / motion / REHO across datasets. This function provides a two-sided test for the null hypothesis that 2 independent samples have identical average (expected) values. It assumes that the populations have identical variances by default.

##### Least squares polynomial fit

python command numpy.polyfit was used to plot the linear regress line in all scatter plot. The solution minimizes the squared error in a polynomial of degree 1.

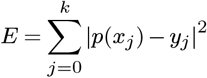

##### Dice Coefficient

Dice Coefficient is a statistical method used to obtain the similarity of two samples. The formula is:

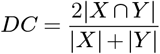

where X, Y are two discrete data sets. We use Dice Coefficient to investigate the similarity of the pooled correlation maps of physiology/motion vs REHO and two connectivity networks, Yeo2011 networks (Yeo et al., 2011) and UK biobank resting-state networks (Shen et al., 2018). Both networks are based on resting-state fMRI. Yeo2011 includes 7and 17-network parcellation estimates by performing clustering on full sample of 1000 subjects. UK Biobank restingstate networks provides major resting-state networks by using group-ICA on fMRI data of more than 4000 UK Biobank participants. ICA was run at two different dimensionalities (25 and 100) and because those components that are not neuronally driven were discarded, 21 and 55 components are included in the final networks. Here, we choose 4.5 for its bordering threshold.

## 3 Results

### REHO vs heart rate (Figure 6 and 7)

Figure 6 shows REHO vs heart rate correlation across subjects, using a significance threshold of p<0.05 for all datasets. Each voxel represents the PCC between the vector of cardiac rate values, and the vector of REHO values. We show a sample scatter plot from a single voxel in the UCLA-control dataset, demonstrating in that particular voxel that subjects with higher heart rate had lower REHO (each point is a single subject). High inverse correlations (p<0.05) are evident across most datasets. Subjects who had lower heart rate tended to have higher REHO. One dataset (InterTVA) shows relatively high positive correlation. Averaging the correlation maps across all 9 datasets (Figure 7(Left)) showed widespread, inverse relationship between heart rate and REHO, with peak values in the temporal lobe (an area of dense vascularization). Pooling all subjects together and performing the heart rate vs REHO correlation with all 412 subjects at once showed similar patterns, with slightly weaker correlation values (Figure 7(right)), however these correlations were highly significant due to the larger sample size with a significance threshold of p-value<0.05.

**Fig. 6.**
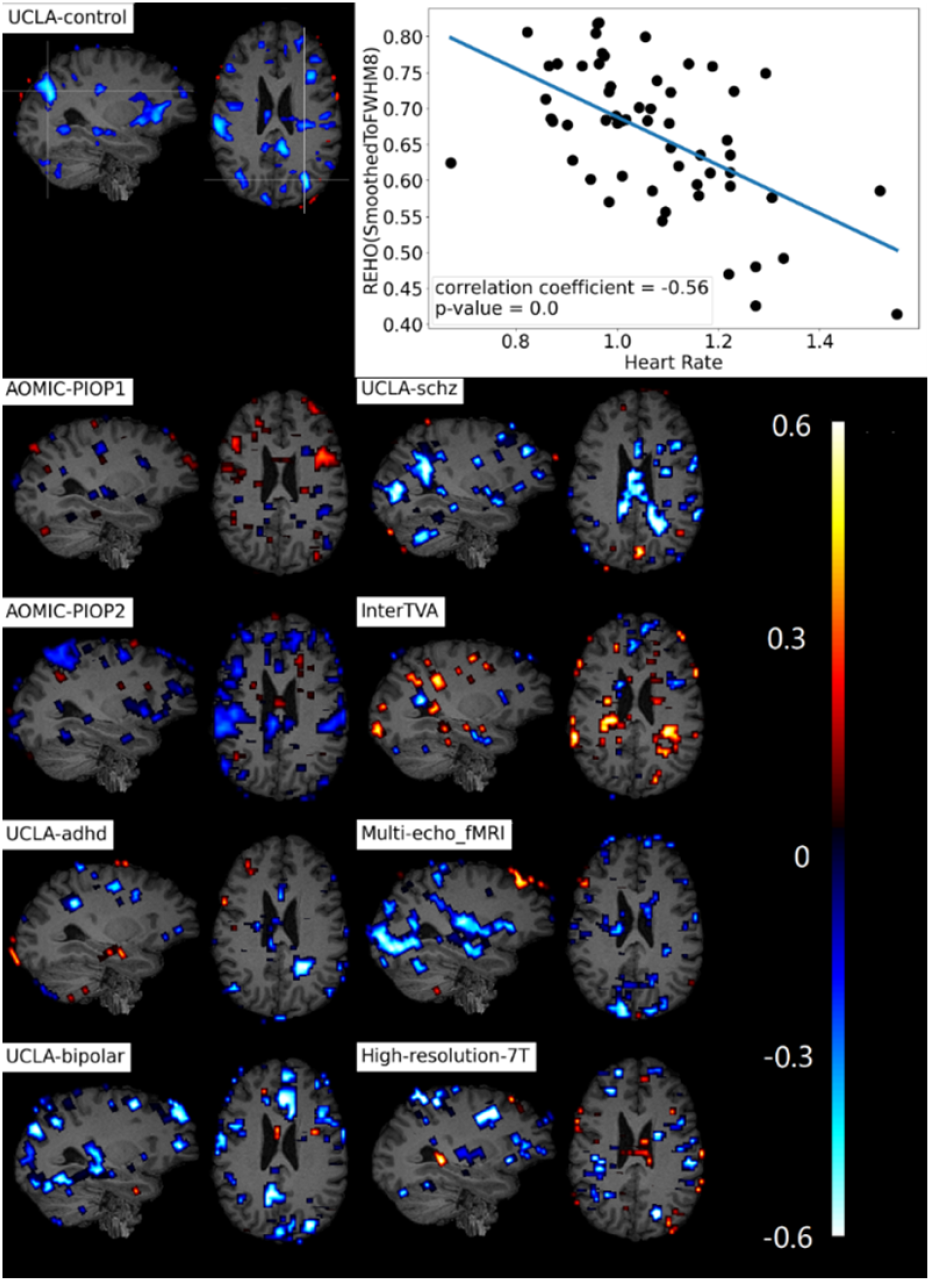
Correlation maps of REHO vs Heart Rate across subjects inter-group heterogeneity. 9 datasets’ correlation maps and a scatter plot showing the correlation in the cross marked voxel of the correlation map of data-set UCLA-control.

**Fig. 7.**
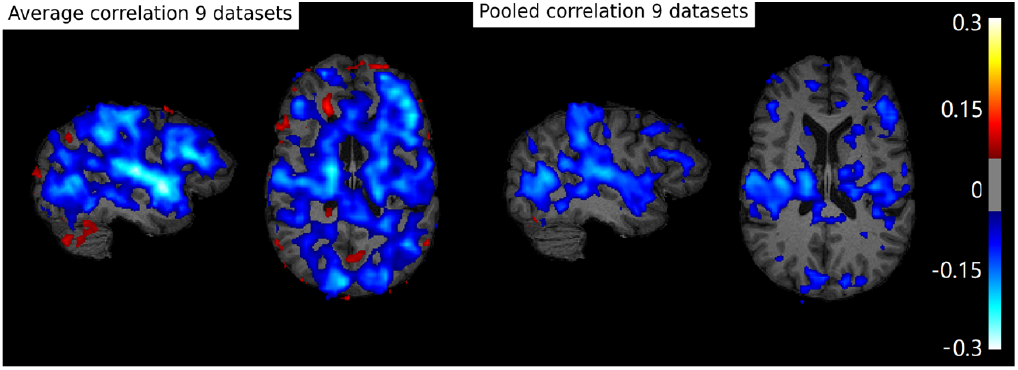
Correlation maps of REHO vs Heart Rate across subjects inter-group heterogeneity. (Left) average correlation map of 9 correlation maps in (Fig.6). (Right) Correlation map calculated by pooling all subjects of 9 datasets together.

### Dynamic REHO vs respiratory rate (Figure 8-10)

Figure 8 shows a correlation map of one subject in dataset UCLA-control and a scatter plot representing one voxel with peak correlation value (−0.96 with p-value=0). The correlation shows the relationship between REHO and respiratory rate across time in each subject. Figure 9 shows averaged correlation values (threshold = ±0.05) of all subjects in each dataset. In 6/9 datasets, we found significant wide spread inverse relationship between REHO and respiratory rate across time, indicating that REHO was increased temporarily during epochs when the subject’s breathing slowed. Correlations were strongest in the temporal lobe (similar with heart rate), parietal lobe and insula regions. Averaging all datasets (Figure 10(left)), with peak correlation value around 0.2, confirmed this finding, as did the pooled analysis with correlation between 412*10 REHO images and 412*10 respiratory feature values. Pooled correlation is under significance threshold p-value<0.05 and correlation value threshold=0.05.

**Fig. 8.**
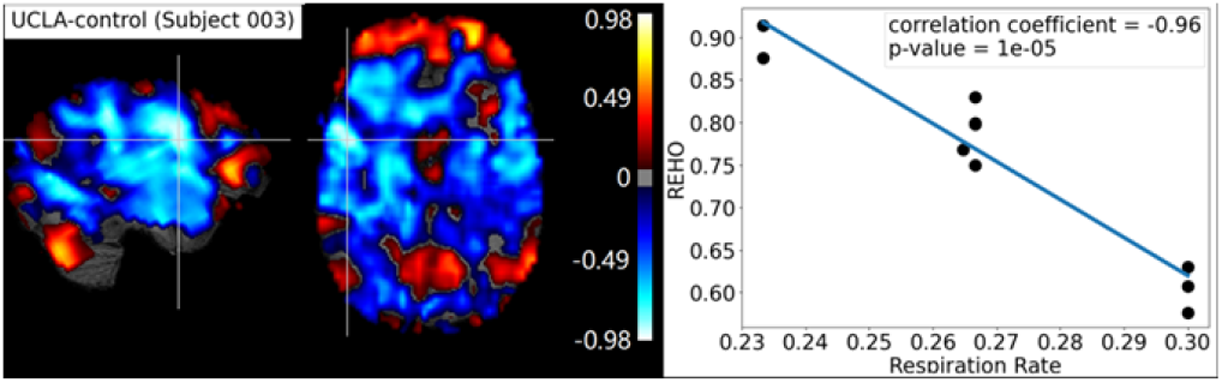
Correlation maps of REHO vs respiratory rate across time inter-subject heterogeneity. 10 slices window correlation map of a subject003 in dataset UCLAcontrol and scatter plot showing the REHO and respiratory rate values in the cross marked voxel.

**Fig. 9.**
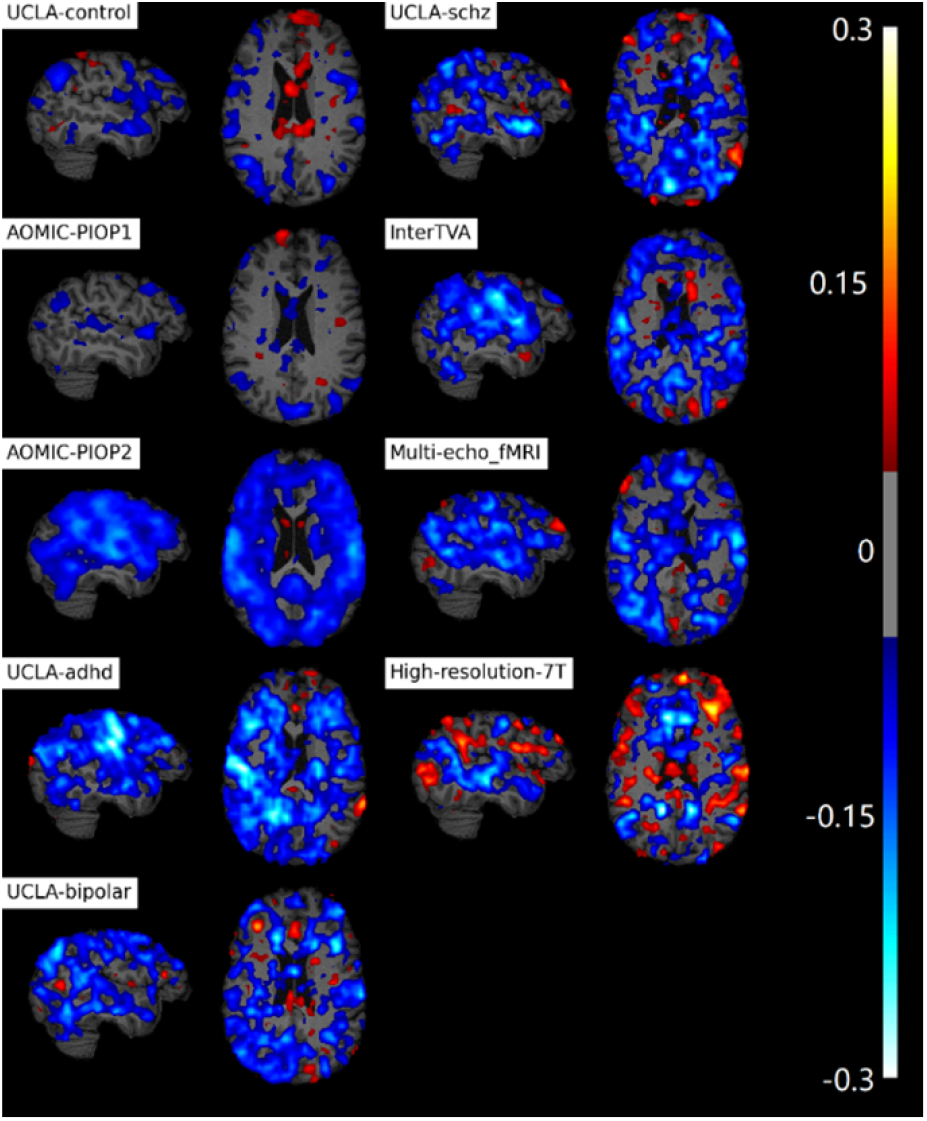
Correlation maps of REHO vs respiratory rate across time inter-subject heterogeneity. Averages of 10 slices window correlation maps of all subjects in 9 datasets.

**Fig. 10.**
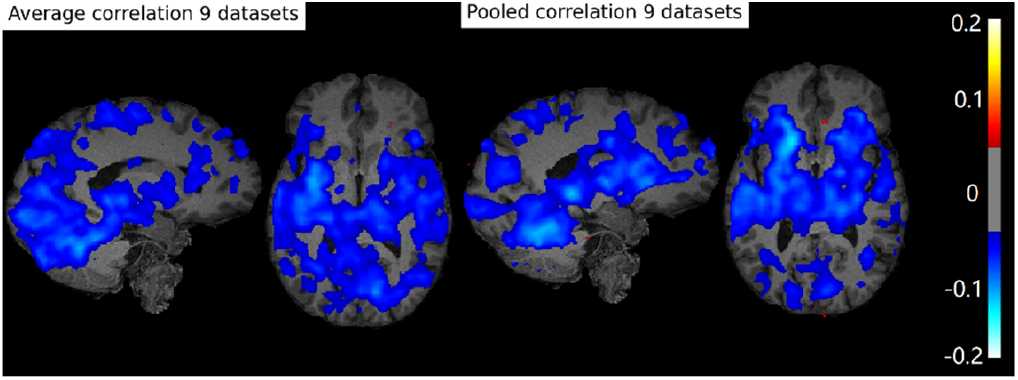
Correlation maps of REHO vs respiratory rate across time inter-subject heterogeneity. (Left) Averaging correlation map of all 9 datasets. (Right) Correlation map calculated by pooling all 9 datasets together.

### REHO vs motion (Figure 11 and 12)

Figure 11 shows correlation map of REHO vs motion of 9 datasets and a scatter plot for one single voxel correlation (0.89 with p-value=0.00012) in dataset ‘high-resolution-7T’. Correlation maps were corresponding to a significance threshold of p-value<0.5 and r-value threshold=0.1. The correlation map of dataset ‘high-resolution-7T’ shows high positive correlation (over 0.8) in parietal lobe and temporal lobe. In this dataset, REHO vs motion revealed that subjects who moved more, tended to have higher REHO values. But for other datasets, there is no clear widespread correlation. 6/9 datasets showed no clear relationship and 2/9 datasets showed an inverse relationship between REHO and motion. Dataset UCLA_bipolar showed relatively strong inverse correlations (around −0.6) in temporal lobe. This resulted in a lack of widespread correlation when averaging correlation maps across all datasets as well as pooling all subjects to produce a single correlation map (significance threshold p-value<0.05) (Figure 12(left)). both the average and pooled correlation maps show positive correlation between REHO and motion (r>0.15) in temporal lobe and edges of brain.

**Fig. 11.**
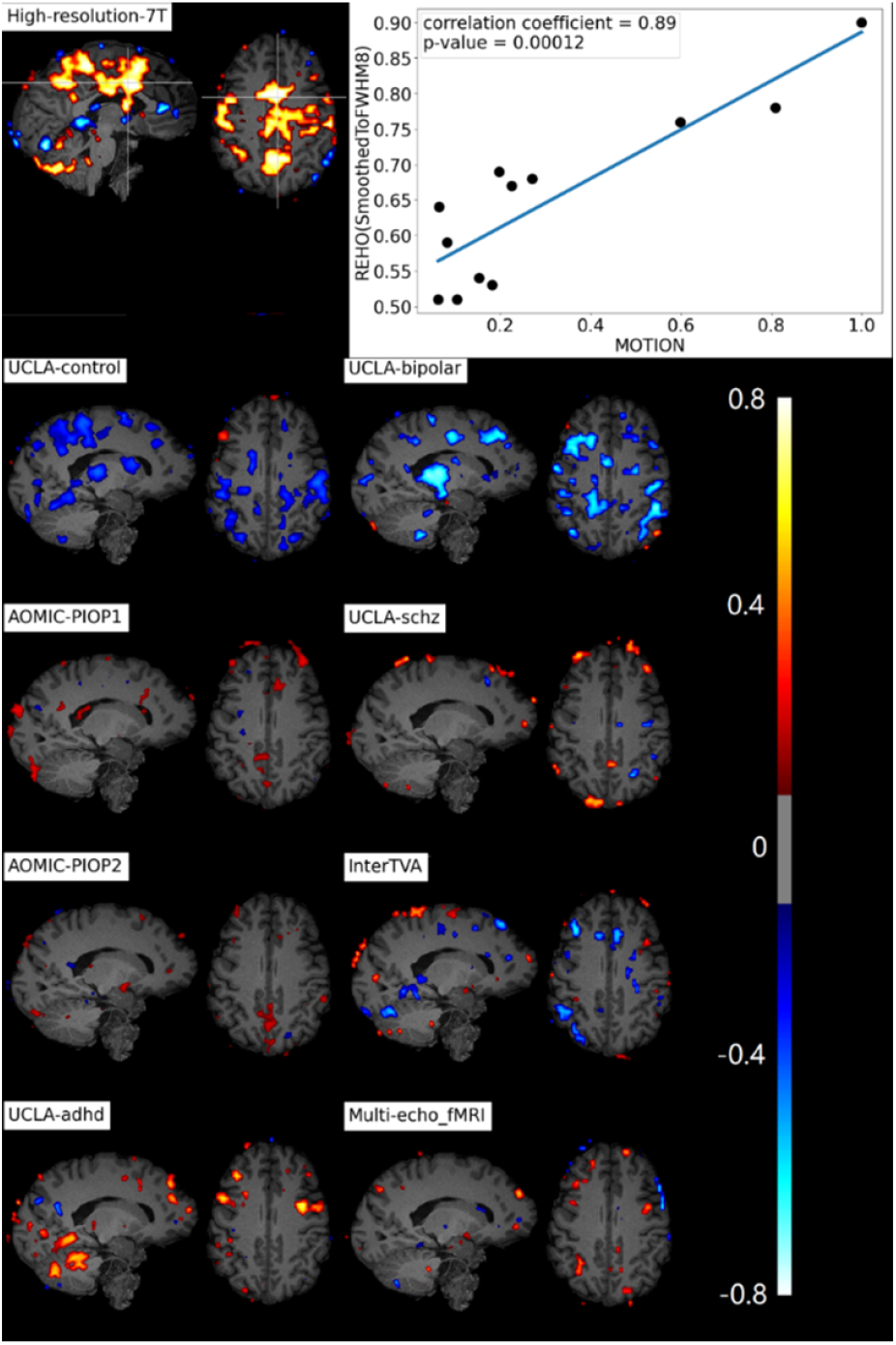
Correlation maps of REHO vs Motion Parameters across subjects intergroup heterogeneity. 9 datasets’ correlation maps and a scatter plot showing the correlation in the cross marked voxel of the correlation map of data-set High-resolution-7T.

**Fig. 12.**
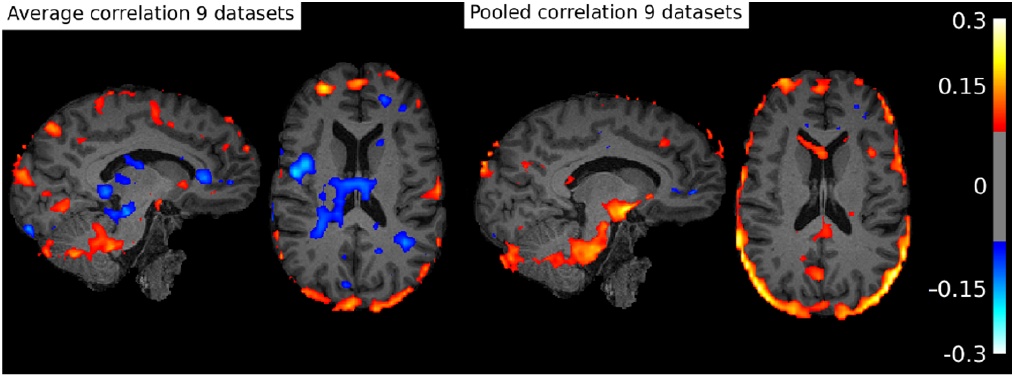
Correlation maps of REHO vs Motion Parameters across subjects inter-group heterogeneity. (Left) average correlation map of 9 correlation maps in (Fig. 11). (Right) Correlation map calculated by pooling all subjects of 9 datasets together.

### Differences in physiological parameters and REHO across groups (Figure 13-16)

We quantified mean heart rate, respiratory rate, and motion across all datasets (Figure 13). As expected, heart rate was close to 60 beats/minute across all datasets, although the schizophrenia group (dataset 5, UCLA-schz) had significantly higher heart rate than other groups. Respiratory rate was around 16 breaths per minute, with dataset (5) UCLA-schz and dataset 8 InterTVA have the highest respiratory rate (Figure 13, center). The difference of mean motion parameter among 9 datasets is more obvious than heart rate and respiratory rate. Dataset (5) UCLA-schz again had the highest value while dataset High-resolution-7T has the lowest value. As expected, all disorder groups (groups 3,4,5) (Figure 13, right) had high motion values, indicating that disorder groups had more difficulty remaining still in the scanner. Other two datasets AOMIC-PIOP1(6) and AOMIC-PIOP2(7) also have relative high motion value, possibly due to the younger average ages.

**Fig. 13.**
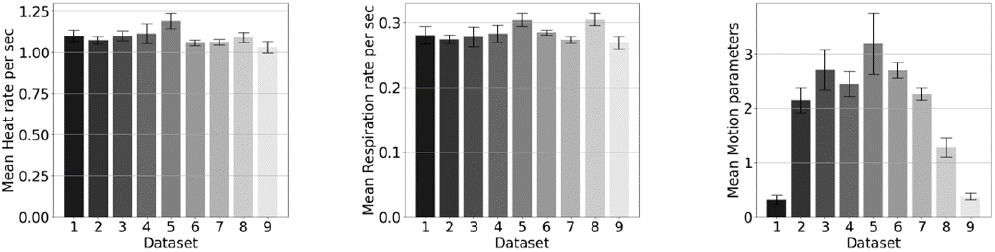
Mean, standard errors and t-test. Datasets are 1: High-resolution-7T; 2: UCLA-control; 3: UCLA-adhd; 4: UCLA-bipolar; 5: UCLA-schz; 6: AOMIC-PIOP1; 7: AOMIC-PIOP2; 8: InterTVA; 9: Multi-echo_fMRI. REHO includes REHO without processing, REHO after FWHM and REHO after FWHM and regression out of Respiratory Rate, Heart Rate and Motion Parameters. Means and standard errors of heart rate, Respiratory Rate, Motion.

**Fig. 14.**
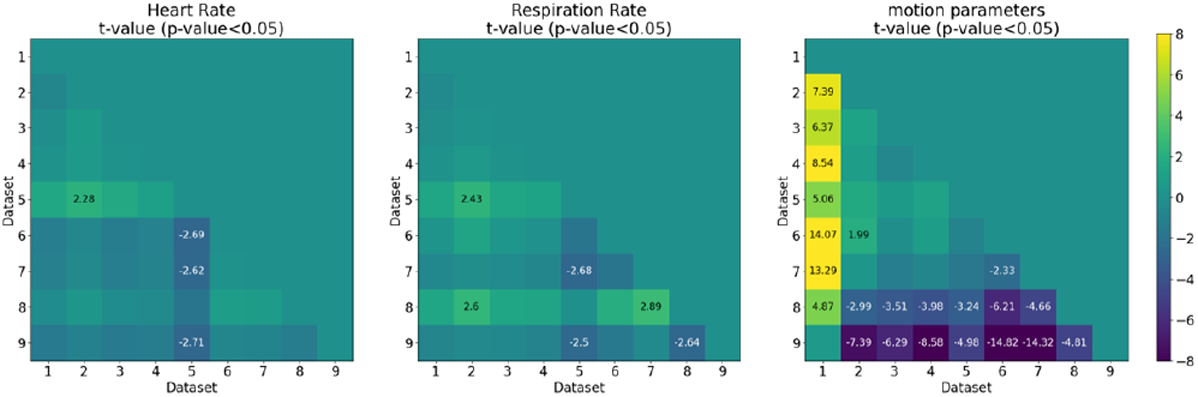
Mean, standard errors and t-test. Same datasets and parameters as described in Fig. 13. p-value and t-value of heart rate, Respiratory Rate, Motion.

In contrast to physiology measures (which were relatively stable across datasets), average REHO (full-brain average) values were highly heterogeneous across datasets, with significant difference in full-brain REHO across nearly all groups (Figure 15 left). The datasets AOMIC_PIOP1 (6) had lowest REHO (0.26), while InterTVA (8) and Multiecho_fMRI (9) had the highest REHO. After applying a ‘BlurToFWHM’, mean REHO values were increased across all datasets (Figure 15 center), and significant differences in REHO were reduced across datasets. After BlurToFWHM, the dataset AOMIC_PIOP1 (6) had the highest REHO. Finally, we regressed out the effects of heart rate, respiratory rate, and motion from the mean REHO across the entire brain (see methods) and recomputed the mean REHO across the subjects in each dataset (Figure 16, right).

**Fig. 15.**
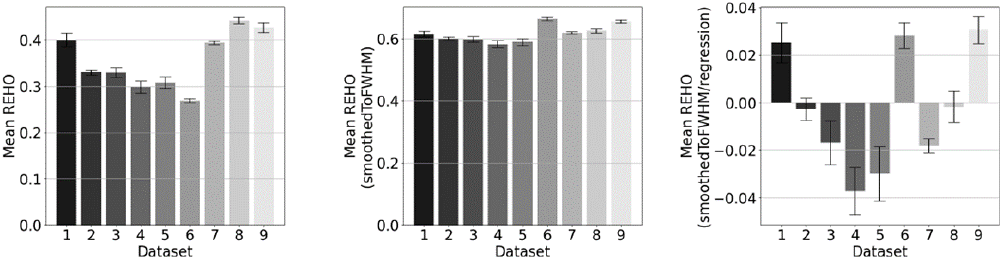
Mean, standard errors and t-test. Same datasets and parameters as described in Fig. 13. Means and standard errors of REHO of 9 datasets.

**Fig. 16.**
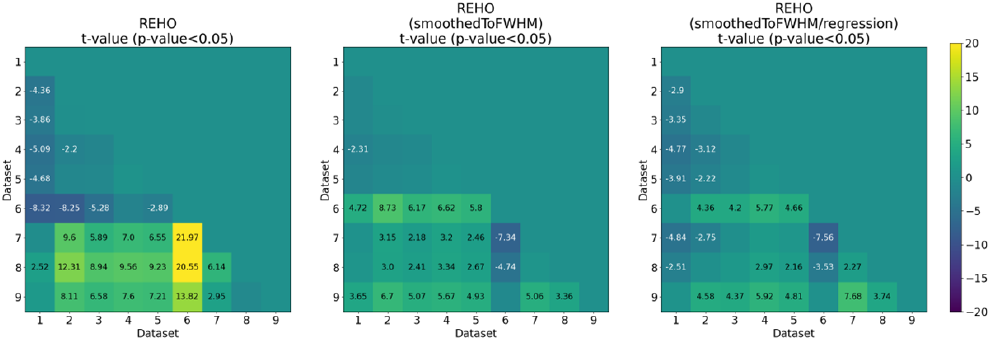
Mean, standard errors and t-test. Same datasets and parameters as described in Fig. 13. p-value and t-value of REHO of 9 datasets

### Dependence of REHO on physiology/motion/FWHM across groups (Figure 17-21)

To examine the relationship between physiological/motion/FWHM parameters and REHO at the group level we plotted mean full-brain REHO as a function of physiological parameter on a 9-point scatter plot (9 datasets). Without performing any correction for FWHM (Figure 17), we find there is no significant correlation with heart rate (p=0.3) and respiratory rate (p=0.9) but motion and FWHM were strongly associated to REHO at the group level; groups with higher motion had lower REHO (r=-0.79, p=0.01) while groups with higher FWHM in the raw data had higher REHO (r=0.78, p=0.01). Figure 18 shows correlation maps where each voxel’s value was obtained from a 9-point correlation between REHO in that voxel (average in each group) and average physiological parameter for each group. The maps were under the threshold at p<0.05. There is no distinct regional correlation for heart rate vs mean REHO or for respiratory rate vs mean REHO. However, nearly the entire brain shows high inverse relationship between REHO and motion (r>0.7).

**Fig. 17.**
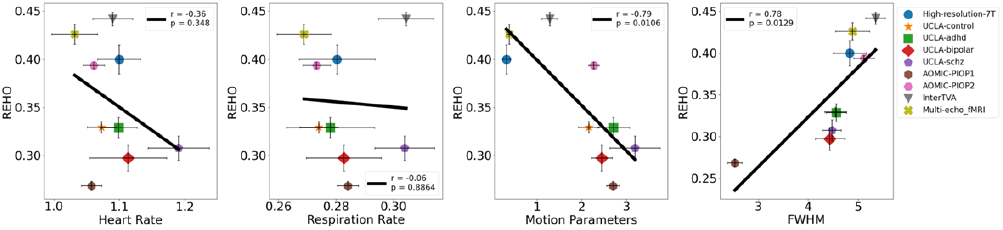
Correlation REHO vs physiological artifact. Scatter plots of REHO (before smoothing and regression out of Respiratory Rate, Heart Rate and Motion Parameters) vs Respiratory Rate, Heart Rate, Motion Parameters and FWHM.

**Fig. 18.**
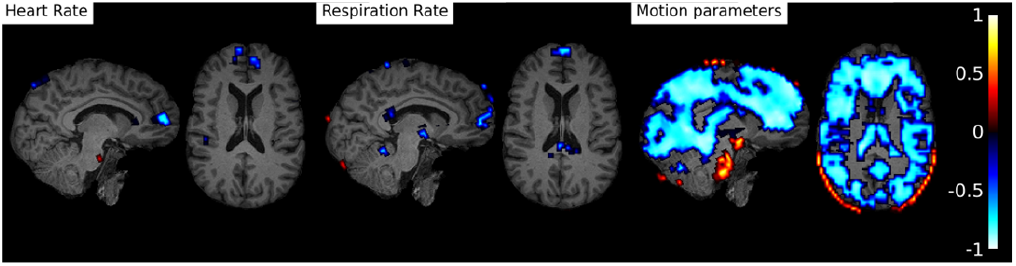
Correlation REHO vs physiological artifact. Correlation maps of Heart Rate, Respiratory Rate, Motion across vs REHO (before smoothing and regression out of Respiratory Rate, Heart Rate and Motion Parameters) across 9 datasets.

After correcting for FWHM using BlurToFWHM (Figure 19), the REHO vs respiratory rate still shows no association at the group level (p=0.55). However, the REHO vs heart rate shows strong inverse correlation (r=-0.72, p=0.03) indicating that differences in raw data FWHM may obscure the heart rate vs REHO correlation, or that smoothing increases contribution of heart rate to REHO. After BlurToFWHM, the strong association between REHO and motion across datasets decreased from r=-0.79 to r=-0.41 with p=0.3, and FWHM also became less associated with REHO (p=0.3) (as expected). Figure 20 shows correlation across groups after BlurToFWHM; significant negative correlation in heart rate vs mean REHO shows in frontal lobe and temporal lobe. There is still no distinct correlation in respiratory rate vs mean REHO. High inverse relationship between mean fullbrain REHO and motion disappeared but in frontal lobe negative correlation remains high (>0.7).

**Fig. 19.**
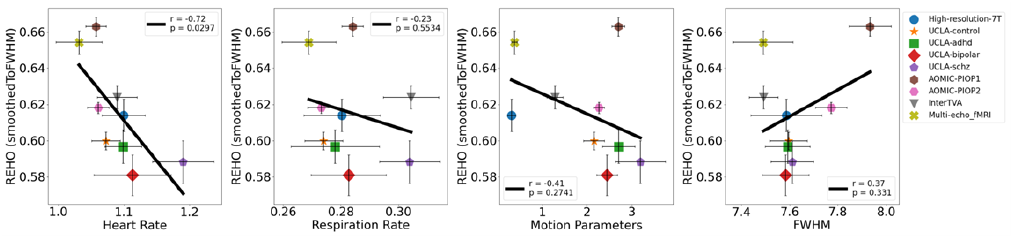
Correlation REHO vs physiological artifact. Scatter plots of REHO (after smoothing and before regression) vs Respiratory Rate, Heart Rate, Motion Parameters and FWHM.

**Fig. 20.**
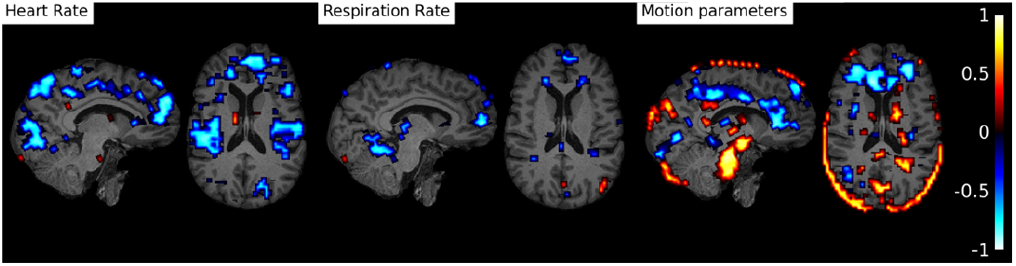
Correlation REHO vs physiological artifact. Correlation maps of Heart Rate, Respiratory Rate, Motion across vs REHO (after smoothing and before regression) across 9 datasets.

**Fig. 21.**
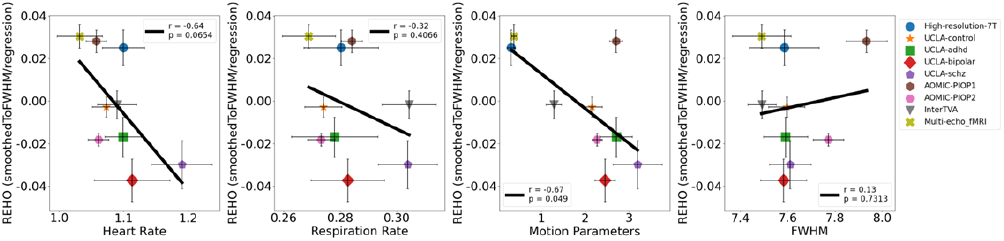
Correlation REHO vs physiological artifact. Scatter plots of REHO (after smoothing and after regression) vs Respiratory Rate, Heart Rate, Motion Parameters and FWHM.

Finally, we examined group-level REHO vs heart rate, respiratory rate, motion, and FWHM after both FWHM correction and regressing out the effects of heart rate/respiratory rate/motion. As expected, the correlation between heart rate and REHO was reduced to r=-0.64, p=0.07 and respiratory rate and FWHM remains not associating with REHO at the group level after these corrections (p=0.32, p=0.73). However, motion vs REHO shows a trend towards an increasing inverse relationship (r=-0.67, p=0.049).

### Dice Coefficient (Figure 22-25)

Finally, Dice Coefficients were calculated to investigate the similarity of the pooled correlation maps (Figure 7(right), Figure 10(right) and Figure 12(right)) with Yeo2011 7-, 17-networks and UK Biobank RSNs ICA-25, ICA-100. Heart Rate correlation map shows high similarity with Yeo2011 7-network #2 (0.81) (Figure 23 left) and Yeo2011 17-network #3 and #4 (over 0.8) (Figure 23 right). At the mean time high similarity with UK Biobank networks shows in ICA-25 #10, #11, #12, #17 (over 0.6) and ICA-100 #2, #6, #20, #22, #30, #32, #34, #35, #41(over 0.6). Figure 25 shows the regions of maximum dice coefficient in color cyan (0.78 in ICA-25, 0.8 in ICA-100). Comparing to Heart Rate, Respiratory Rate correlation map has less similarities in Yeo2011 networks. Dice coefficients higher than 0.6 show in Yeo2011 7-network #2, #4 and Yeo200 17-network #2, #4, #7, #8. On the contrast, Respiratory Rate correlation map has higher dice coefficients with UK Biobank networks, 0.8 in ICA-25 #18 and 0.89 in ICA-100 #38. Figure 25 shows the regions in color magenta. Motion pooled correlation map has relatively low dice coefficients with Yeo2011 networks (under 0.3) and high similarity (0.69) with UK Biobank network ICA-100 #17 shown in Figure 25 in color yellow.

**Fig. 22.**
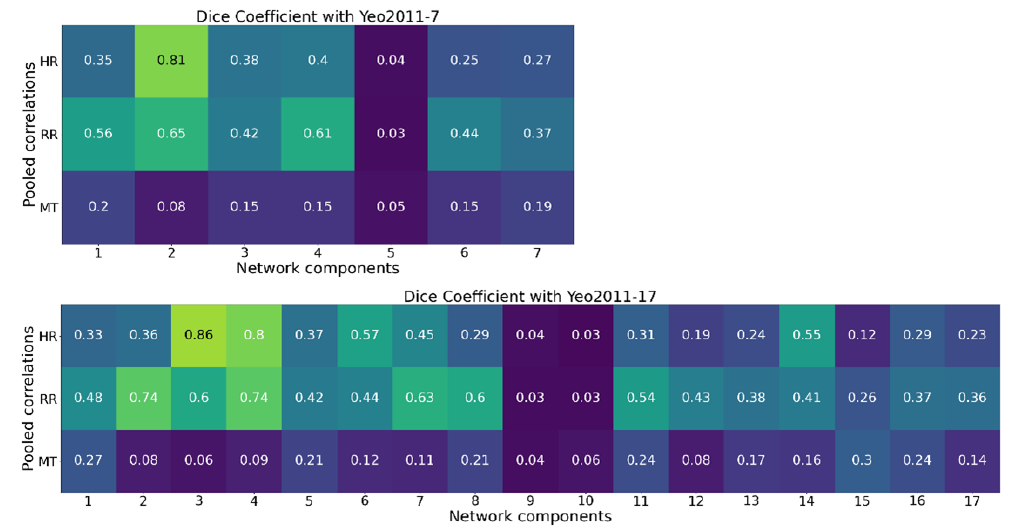
Dice Coefficient. Dice Coefficient between pooled correlations (shown in Figure 7(right), Figure 10(right) and Figure 12(right)) and Yeo2011 network (7 components and 17 components).

**Fig. 23.**
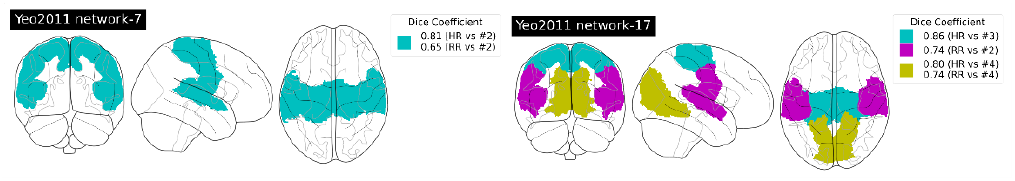
Dice Coefficient. Map of the components with highest Dice Coefficient.

**Fig. 24.**
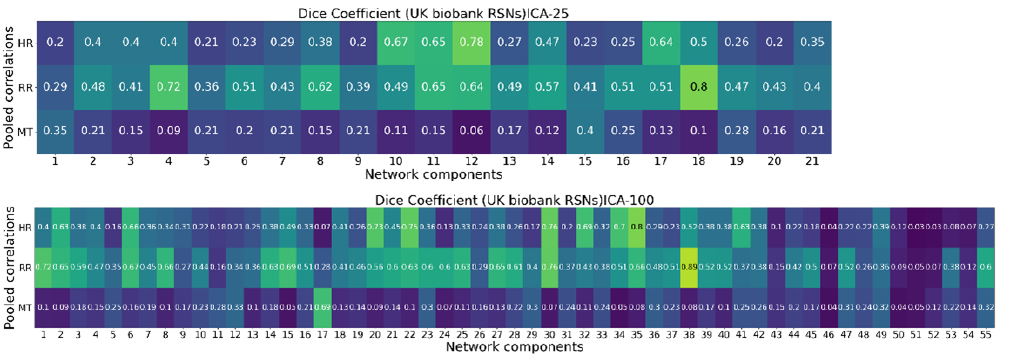
Dice Coefficient. Dice Coefficient between pooled correlations (Figure 7(right), Figure 10(right) and Figure 12(right)) and UK Biobank RSNs (21 components and 55 components).

**Fig. 25.**
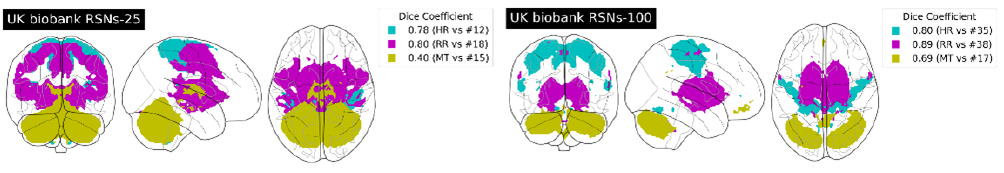
Dice Coefficient. Map of the components with highest Dice Coefficient.

## Discussions

### Summary

We analyzed 9 separate datasets from Open-Neuro for systematic effects of physiology, motion, and FWHM on REHO estimates. We find strong correlations between REHO and physiology/motion at both the single-subject level and group level, leading us to believe that many of the REHO effects reported in the literature are caused by group differences in physiology/motion, rather than intrinsic brain activity. In particular, patient populations studied with REHO such as schizophrenia have strong systematic differences in physiology (increased heart rate) relative to control groups which could lead to much greater BOLD signal differences than might be expected from differences in intrinsic brain activity. Both heart rate and respiratory rate appear to have an inverse relationship with REHO.

### Heart rate

inverse association occurs in temporal lobe with both inter-subject analysis (Figure 7 (left), (right)) and intergroup analysis (Figure 20) indicating subjects or groups with slower heartbeats have higher REHO, particularly in areas of dense vascularization with many veins and arteries. Changed REHO implies changed hemodynamic response (Zang et al., 2004) while cardiac pulsatility generates small movements in brain tissue as well as inflow effects in and around vessels (Dagli et al., 1999) and changes of heart rate are related to changes in blood flow and blood oxygenation. Heart rate can significantly alter fMRI of resting-state (Chang et al., 2009) especially in the areas close to large arteries and draining veins or edges of the brain, lateral ventricals and sulci (Dagli et al., 1999). Temporal lobe is an area of dense vascularization which explains why a strong association between heart rate and REHO is observed in this area. One should be cautious when claiming that increased or decreased REHO in temporal lobe represents neurodegeneration or altered processing.

### Implications for dynamic functional connectivity

recently, dynamic functional connectivity has received increased attention, as it is thought to provide a mean for analyzing changes in ‘brain state’ across time. Some studies have even examined differences in ‘dwell state’ between healthy controls and patient population such as schizophrenia (Damaraju et al., 2014). Here, by breaking up the time series into 10 ‘segments’ we find in all 9 datasets an inverse association between respiratory rate and REHO across time – when a subject breathes more slowly, their REHO increases. This effect could confound studies of DFC, as removing effects of respiratory rate from the BOLD signal is notoriously difficult.

### Motion

motion artifacts have been recognized as potential confound for studies of RSFC for many years already(Power et al., 2012), however we are not aware of any studies explicitly looking at effects of motion on REHO. We show in Figure 21 that groups with increased motion levels have reduced REHO, in fact, before correcting the raw data using BlurToFWHM, motion explains 67% of REHO variance across groups. This is highly problematic for studies on patient populations (bipolar, schizophrenia, adhd) as these populations are well known to move more in the scanner (which we also show in Figure 13). Interestingly, the inverse motion vs REHO correlation was decreased after BlurToFWHM, this could indicate that increased motion reduces the ‘smoothness’ of the data (and, consequently, REHO estimates), thus, smoothing the data may help reduce effects of motion on rs-FMRI measures. Some evidence supports this (Scheinost et al., 2014) showed smoothing all images to a uniform level across the sample, is an effective way to reduce motion-related confounds in functional connectivity studies.

### Inter-group differences

smoothing reduced the differences in mean full-brain REHO among different datasets which means smoothing fMRI data to a certain FWHM is a necessary step for performing inter-group REHO analysis. In fact, we show here that differences in REHO across groups are likely to be due to differences in data smoothness, if BlurToFWHM is not used prior to REHO calculation. On the other hand, smoothing may increase the effects of physiology on REHO inter-groups. Smoothing increased the correlation of heart rate vs mean full-brain REHO in group-based correlation (Figure 17, 19 heart rate). This may be caused by the same reason as that spatially smoothing high resolution data might reduce time series SNR compared to direct acquisition at lower spatial resolution (Triantafyllou et al., 2006).

Paradoxically, regressing motion/heart rate/respiratory rate from the data increased the correlation between motion and REHO at the group level. This can be explained by analyzing the low correlation of motion vs full-brain smoothed REHO firstly. After smoothing, correlation of motion vs REHO turns to positive in temporal lobe and keep being negative in frontal lobe comparing to the whole brain negative correlation of motion vs raw REHO (Figure 18, 20). Due to the coexisting negative and positive correlations, the scatter plot in Figure 19 shows great decrease in the correlation of the full-brain average REHO vs motion. We can see that large proportion of negative correlations returns but is weaker than the association of motion vs raw REHO (Figure 17, 21). As a result, regressing motion from mean REHO at the group level leads to slight increase correlation between REHO and motion (Figure 19, 21).

We show that regressing out the effects of physiology and motion reduces the differences in REHO across datasets (Figure 16), indicating that group difference in REHO is highly susceptible to differences in physiology/motion. We should also note that REHO differences between disease and control in the UCLA cohorts were weaker than REHO differences between UCLA control and other control groups (from other datasets), even after applying BlurToFWHM and regressing out physiology/motion parameters. This indicates that it is probably meaningless to compare REHO across studies where the scanning environment and fMRI sequence parameters were not tightly controlled and uniform. Finally, the dice coefficients shown in Figure 22-25 indicates that correlation maps of Heart Rate, Respiratory Rate and Motion vs REHO are very similar with resting-state networks in certain regions, which proved again that functional connectivity of those regions in resting-state are probably due to physiology/motion artifacts.

